# A physical model for M1-mediated influenza A virus assembly

**DOI:** 10.1101/2024.06.01.594783

**Authors:** Julia Peukes, Serge Dmitrieff, François J. Nédélec, John A. G. Briggs

## Abstract

Influenza A virus particles assemble at the plasma membrane of infected cells. During assembly all components of the virus come together in a coordinated manner to deform the membrane into a protrusion eventually forming a new, membrane-enveloped virus. Here we integrate recent molecular insights of this process, particularly concerning the structure of the matrix protein 1 (M1), within a theoretical framework describing the mechanics of virus assembly. Our model describes M1 polymerization and membrane protrusion formation, explaining why it is efficient for M1 to form long strands assembling into helices in filamentous virions. Eventually, we find how the architecture of M1 helices is controlled by physical properties of viral proteins and the host cell membrane. Finally, by considering the growth force and speed of viral filaments, we propose that the helical geometry of M1 strands might have evolved to optimize for fast and efficient virus assembly and growth.

**Significance:** Influenza A virus remains a major threat to public health. Its most abundant viral protein, matrix protein 1 (M1), forms an endoskeleton underneath the viral membrane, but how this endoskeleton contributes to the virus’ lifecycle is poorly understood. Combining cryo-electron tomography data and structural data with theoretical predictions, we explain how the energetically favorable polymerization of M1 into helical strands mediates the membrane deformations that permit the virus to exit infected cells. Our analysis of M1’s variable architecture provides insights into adaptive strategies of the virus for efficient growth under variable local conditions. The quantitative framework developed in this study could be extrapolated to other enveloped viruses and generally applied to protein-driven membrane deformations.

## Introduction

Influenza A virus (IAV) represents a major threat to global health, infecting 100,000s of people per year during seasonal epidemics and occasional pandemics (1). Influenza virions are enveloped particles with variable morphologies that range from small spheres to filaments that can be several micrometres long (2, 3). The filamentous form is observed in the context of human infections e.g. within lung tissues of infected individuals (4). The viral envelope of assembled influenza virions is densely decorated by the glycoproteins hemagglutinin (HA) and neuraminidase (NA) and contains low copy numbers of the ion channel matrix protein 2 (M2) (5–8). The inside of the lipid bilayer is coated by an endoskeleton formed from the matrix protein 1 (M1) (9, 10). The segmented viral genome, packaged within viral ribonuleoproteins (vRNPs), is typically located at the front tip of the virus (10, 11).

During an infection, new influenza virions assemble at the plasma membrane of infected cells (12). Virus assembly is driven by interactions of HA, NA, M1, M2 and the vRNPs with each other and with the plasma membrane (13, 14). Details of how this process is coordinated have remained elusive. Expression of HA or NA together with M1 is sufficient to assemble filamentous particles that closely resemble virions while in particular expression of HA alone gives rise to pleiomorphoic, non-filamentous particles (15).

M1 has been considered a primary mediator of virus assembly: Structural and genetic studies of virus assembly and viral proteins have shown that specific, single point mutations in M1 can impact virus morphology (16–18). Within virions, M1 forms a tight protein meshwork directly underneath the viral envelope that may play a key role in virus assembly by interacting with the membrane and all other viral components.

To understand how M1 polymerization mediates virus assembly, we recently acquired and analyzed cryo-electron tomography (cryoET) data of influenza A/Hong Kong/1/1968 (H3N2) (hereafter HK68) virions budding from cells (19). Using subtomogram averaging we studied the *in situ* organization and structure of M1 and found that M1 forms linear polymers underneath the membrane. From this *in situ* structure in combination with high-resolution full-length *in vitro* M1 structures from us and others (19, 20), we know that M1 forms linear strands via a hydrophobic interface between N- and C-terminal domains from neighbouring M1 monomers. Inside virions, linear M1 polymers arrange as multiple parallel helical strands. The number (1–7) and the helical handedness of those M1 strands can vary between individual virions (19). The function of this variability or a potential role in the context of virus assembly remains unclear.

Assembly and release of enveloped influenza virus particles requires reorganization of the plasma membrane. Understanding membrane deformation is key to fully understand and describe virus assembly, but membrane parameters that govern membrane behavior are difficult to accurately measure inside complex systems such as virus infected cells. Theoretical modeling of membrane deformation processes can provide insight into the complex interplay of membrane parameters, linking molecular mechanisms on the protein level with the observed membrane behavior on the cellular level. Since the pioneering work by Helfrich (21), membranes are often modeled as elastic surfaces in which rigidity and tension oppose membrane deformation. This approach allows ones to predict the shape of a membrane under constraints, such as point forces or pressure difference. Membrane-associated proteins are represented either by modifying some membrane parameters, or as an external field, depending on the biological context. For yeast endocytosis, for instance, the protein coat was included by choosing a much larger effective membrane rigidity, compared to the membrane alone

(22). In filopodia a linear bundle of actin filaments extrudes a membrane tube by polymerizing at its tip; actin polymerization can be seen as a force applied to the tip (23). Filopodia formation, exocytosis and endocytosis have been modelled extensively, but few authors have investigated the mechanics of membrane deformation by viral proteins (24, 25).

During assembly of filamentous influenza virions, the cellular plasma membrane is deformed into regular tube-shaped particles – resembling filopodia – by viral proteins interacting with the membrane. We wondered if one of the influenza virus proteins takes the dominant role in force generation for membrane deformation during virus assembly. M1 is a possible candidate since M1 is the most abundant protein in the virus (26, 27) and since M1 is essential for the formation of long protrusions during the assembly of filamentous influenza virus particles (15–17).

Here we introduce a mathematical model in which M1 polymerization provides the driving force to elongate filamentous virions, overcoming membrane tension and rigidity. We further analyze existing cryoET data, trying to identify the polymerization direction of M1. From this insight, and recent structural details on M1 monomers and polymers dimensions, we develop a model of M1 polymerization, recapitulating the observed properties and variability of M1 within virions. Finally, adding previously reported values of membrane properties, we model the growth of filamentous virions.

## Materials and Methods

### Analysis of virus and M1 directionality

Sample preparation, data collection and tomogram reconstruction of the data analyzed here have been described in (19). Briefly, samples that have been analyzed here are Influenza A/Hong Kong/1/1968 virions that are produced from MDCK cells grown directly on cryoEM grids (QF AU200 R 2/2, Quantifoil, Germany), infected at a multiplicity of infection <1, and observed until a cytopathic effect was visible. Cells and viruses on grids were then frozen using a Leica GP2 plunger. Tomograms were collected on a Titan Krios with a K2 detector (Gatan) and an energy filter (20eV slit width) using a dose-symmetric bidirectional tilt scheme, with a tilt range of -60 to 60 and a tilt increment of 3 and a total dose of 120-150 electrons/Å^2^.

To understand the growth direction of virions, we identified virions from the tomographic dataset where the vRNP-containing virus tip (defining the front end of the virus) or the remaining connection of the virus to the cell (defining the rear end of the virus), or both, were visible in the tomogram (Fig. 1A). We also identified virions where it was possible to identify the remaining connection of the virus to the cell, and therefore the growth direction, from the medium magnification maps that are collected prior to tomogram acquisition. In parallel we performed per-virus subtomogram averaging of M1 for each virus for which the growth direction could be determined as described in (19). By displaying the obtained subtomogram average for each virus back into the respective tomograms using the positions and orientations obtained from subtomograms averaging we could then compare the orientation of M1 to the respective growth direction of the virus (Fig. 1A).

**Figure 1.**
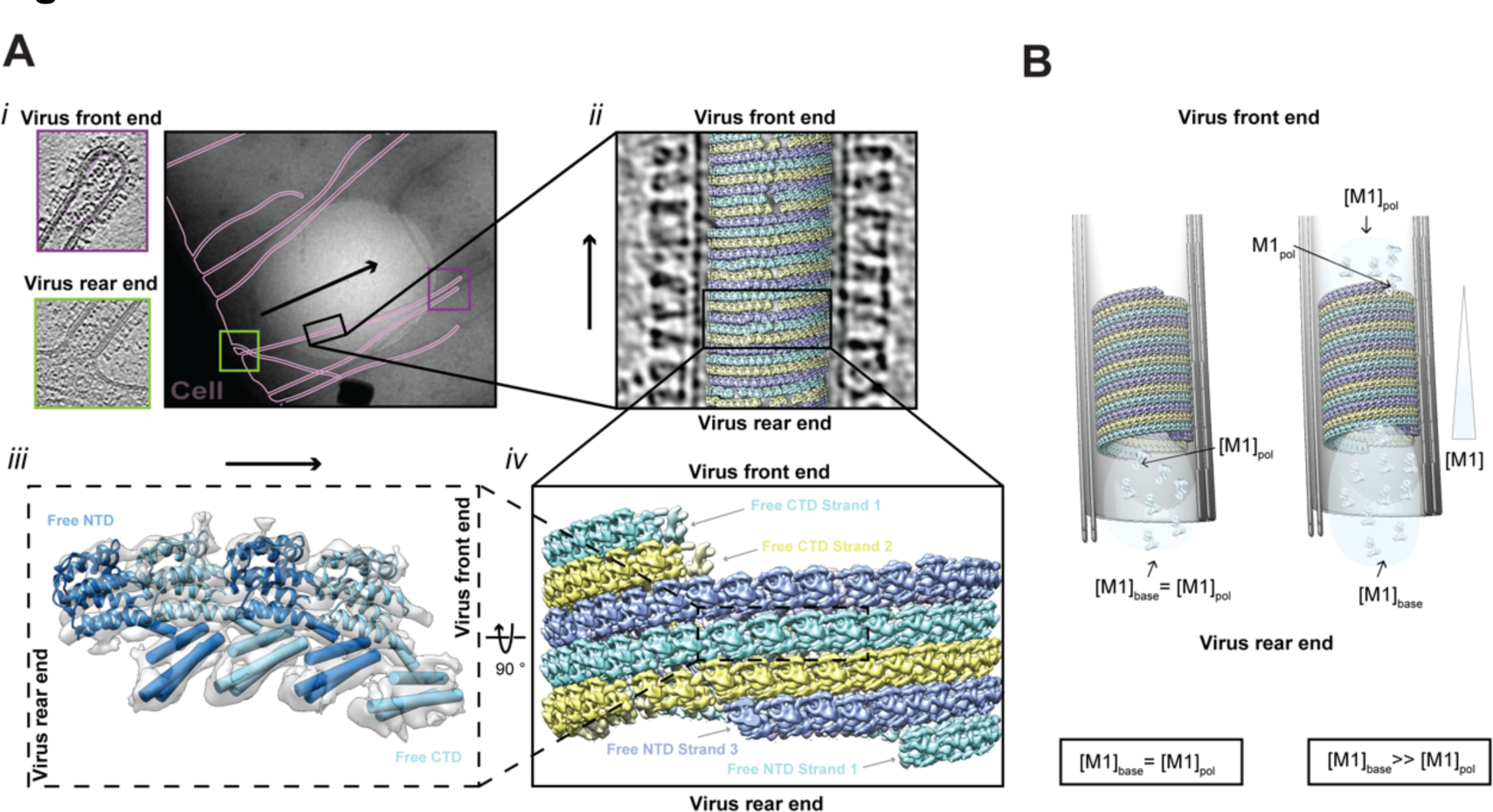
Directionality of virus M1 polymerization. A) Overview of how the direction of virus growth and orientation of M1 are defined and were determined. i) CryoEM image of influenza virions budding from cells – with the outlines of membrane marked in pink. The virus front and rear of one exemplary virus are marked by purple and green boxes respectively. The corresponding purple and green frames show representative tomogram slices of a virus front tip frames, recognizable by the presence of the viral RNPs, and the rear end of virus filament where it is connected to the host cell. ii) Tomogram slice of an influenza A virus filament. A 3D reconstruction of M1 is placed onto the positions of M1 identified by subtomogram averaging of M1. iii) In situ 3D reconstruction of M1 with a model of the M1 NTD fitted and cylinders fitted into the M1 CTD density (EMDB-11077 (19), M1 NTD: PDB 1EA3 (40)) indicating the two distinct ends of an M1 oligomer: Free NTD and Free CTD end. iv) 3D arrangement and directionality of M1 inside the influenza viruses. B) M1 concentration [M1] if M1 polymerizes at the virus base (left) or at the growing virus tip (right).

### Bending rigidity of a M1 strand

To estimate the bending rigidity of a linear strand, we use the formula for the bending rigidity of a slender beam. Calling Y the Young modulus of the material and I the second moment of area of the strand, its bending rigidity should be *Γ_u_= Y I*. For simplicity, we assume strands to have a rectangular cross-section of height *h* and width w, leading to *I = (h^3^w + w^3^h)/12*.

The typical scale of Y for structural proteins is the GPa, leading to *Γ_u_* ∼ 57 10^-27^ J.m. This is our upper estimate, which could be reached if the protein was a single ordered domain. For the lower estimate, we assume on the contrary that the CTD is disjointed and does not participate significantly to the strand bending rigidity; in this case one should take *h* to be only the size of the NTD domain, i.e. 2.5nm, leading to *Γ_u_* ∼ 14 10^-27^ J m.

### Optimal radius of a filamentous virion

We start from equation 1 and assume the filamentous virion to be straight, i.e. the curvature along *v* to be zero. The M1 polymer deformation energy per unit length is thus:

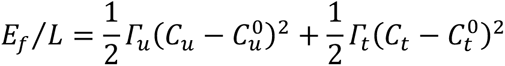

We can use the Frenet-Serret formula to express *C_u_* and *C_t_* as a function of *R* and *b_n_* : *C_u_ = R^2^ / R^2^+b_n_^2^* and *C_t_ = b_n_ / R^2^ + b_n_^2^*. If the helices in the filament are densely packed, as it appears to be the case, experimentally, we can use equations 4 and 5 to express *b_n_* as a function of *R* and *n*. For simplicity, we assume here *Γ_t_*= *Γ_u_* ; relaxing this hypothesis does not alter the qualitative predictions but would change quantitatively the energy minimum. We can then derive *E_f_/L* as a function of R ; the derivative *∂_R_ E_f_ /L=0* should be zero when the energy is at a minimum. We then assume *R* to be close to *1/C<u>_u_</u>^0^* (the radius of curvature of M1 strands along *u*), by writing *R=1/C<u>_u_</u>^0^(1+ε)*, with *ε* small. We then solve *∂_R_ E_f_ /L=0* with respect to *ε*, at second order in *ε*. Eventually, we assume *nh/2π* to be much smaller than *1/C<u>_u_</u>^0^* (as we know from experiments) and we expand to second order in *n h C<u>_u_</u>^0^/2π*, to find:

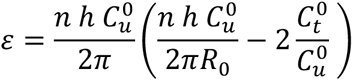

Because of evolution, we expect C_t_^0^ to be close to the torsion observed experimentally, in which case we can write *C_t_^0^∼mhC_u_^0^/2πR_0_^2^*, where *m* is a number expectedly close to 3, the average number of helix starts observed experimentally, yielding 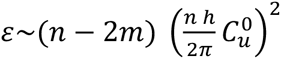.

Therefore, we expect the radius *R* to be *R_0_* plus a small correction, of order (*n h C_u_^0^/2π)^2^*.

## Results

### M1 assembles polarized filaments with a defined directionality

M1 forms a polar polymeric strand with a helical arrangement underneath the membrane of assembling virions. The polarity of the strand can be defined based on the characteristic features of each of the two polymer ends: one end exhibits an unbound M1 NTD and the other end an unbound M1 CTD (Fig. 1A). Here we set out to determine how the polarity of the M1 strand relates to the direction of growth of the filamentous virion. To address this issue, we reanalysed our previously published cryoET data of influenza virions obtained from cells producing virions directly onto EM grids. We identified a subset of virions where the direction of virus growth could be unambiguously determined based on the presence of the viral RNPs at one end of the filament or the location of the producer cell at the other end of the filament. For those 11 virions, we compared the direction of virion growth with the polarity of the M1 strands within the virions. We found for 9 out of 11 virions that the unbound NTD of the M1 polymer was at the base of the virion facing the cytosol (Suppl. Fig. 1). In 8 of those, M1 was arranged into right-handed helical arrays with a variable number of helix starts (1 to 6) while in 1 of those 9, M1 formed a left handed helix. The low number of left handed M1 helices in this subset is representative of the low occurrence of left handed M1 polymers in the full dataset (19). In the remaining 2 out of 11 virions, both of which are right-handed, we found that the M1 orientation was inverted with the free M1 CTD facing the cell body. Those 2 virions also displayed an additional, ordered protein layer inside of M1 (Suppl. Fig. 1). This reflects the frequency of that protein layer in the full data set (13 of 62 virions). Similar looking internal protein layers have been previously observed inside influenza virion (referred to as ‘multilayered coil’ in Fig 1 in (10)) but the identity and function of this protein layer remains to be investigated further.

Thus, in most virions analyzed here and in all virions without an additional inner protein layer, the free-NTD end of the M1 strands faces the cytosol. While it may be natural to assume that assembly of M1 proceeds at the base of the filament facing the cytosol (onto the free NTD), structural data alone cannot rule out the possibility that M1 polymerization could take place at the tip of the filament (onto the free CTD), or at both ends.

### What is the directionality of M1 polymerization?

We here consider the hypothesis that M1 polymerization takes place at the tip of the virion, analogous to polymerization of actin at the tip of filopodia (28). For this to occur M1 monomers must actively or passively travel through the filament to reach its tip. Given the absence of any obvious or previously described filament or other structural feature in the viroplasm that could act in active transport, we consider only passive diffusion. In order for a filamentous virion to grow with a speed *v*, the flux of M1 monomers in the filament towards the tip should be *j* = *v ρ*⁄*πR*^2^ , where *ρ∼*12.5 nm^-1^ is the number of M1 monomers per unit virion length, and *R∼*16 nm is the inner tube radius. Polymerization of M1 at the tip would locally deplete M1 monomers, creating a difference of concentration *Δ*[M1] between the tip and base of the filament (Fig. 1B). The diffusive flux of M1 protein in the filament can be estimated as *j* = *D Δ*[M1]⁄*L_virus_*, with *D* the M1 monomer diffusion coefficient and *L_virus_* the filament length. Therefore, the passive movements of M1 monomers would be fast enough to sustain filament growth if the concentration difference is at least *Δ*[M1] = *vρ L_virus_*⁄*D πR*^2^. We estimate *D*∼10 µm^2^s^-1^, typical for proteins of the size of M1 (29, 30). For a conservative estimate of growth rate of *v*=1 µm/h, and a filament length of *L_virus_*=1 µm, we find *Δ*[M1]∼1 µM. Thus, with unhindered diffusion in the filament, the observed polymerization rates require a cytoplasmic concentration of M1 larger than 1 µM. For longer virions the concentration would need to be higher, proportionally to the virion length, unless their growth is slower (>10 µm virions are frequently observed by us and others (18, 31, 32)). Since cellular M1 concentrations of up to 10 µM have been reported in a transfection based system (33), this scenario is theoretically possible. However, we consider more likely that M1 polymerizes at the filament base which is not hindered by the viral genome and where polymerization has much less stringent requirements on M1 concentration.

### A physical model for influenza virus filament protrusion

We next asked whether polymerization of M1 could provide the energy for filamentous virus protrusion, and whether this assumption can be consistent with the observed range of strand parameters and filament diameters. To answer this question, we consider a physical model in which the membrane, coated by the glycoproteins HA and NA (Fig. 2A), behaves like an elastic surface of elastic modulus *K_m_*, spontaneous curvature R_0_, and tension σ. Polymerized M1 is represented as a ribbon with three principal stiffnesses *Γ_u_*, *Γ_v_*, *Γ_t_* and their associated spontaneous curvatures *C_u_^0^*, *C_v_^0^*, *C_t_^0^*, with *C_t_^0^* being better understood as a spontaneous torsion (Fig. 2B). These parameters are summarized in Table 1. We will assume that these quantities are constant all along M1 strands. Calling *δL* the M1 strand length gained by adding one monomer of length *a* (table 1), and *δG* the change in free energy associated with this extension, the polymerization force is f=*δG*/*δL*.

**Figure 2.**
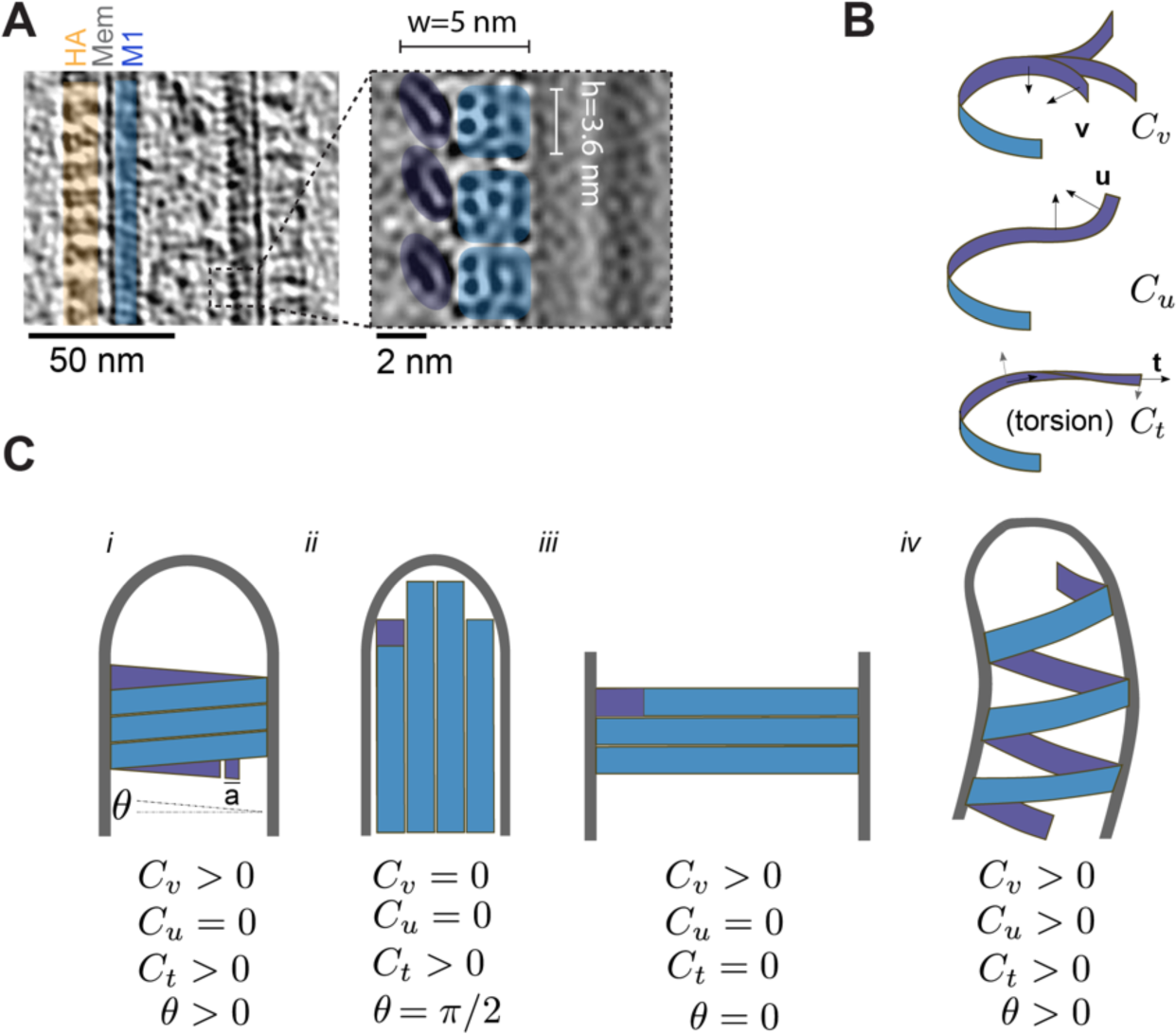
M1 dimensions and mechanical properties predict different scenarios for M1 polymer architecture and virus filament shape. A) Left: CryoET slice of a section of an influenza A virus. The different components: HA (hemagglutinin), Mem (Membrane) and M1 are marked. Right: XZ-orthoslice through the 3D reconstruction of three neighbouring M1 strands. The dimensions of M1 monomers measured from this reconstruction are indicated. Adapted from (19). B) Mechanical properties of polymerized M1 represented as a filament with three principal spontaneous curvatures *C_u_^0^*, *C_v_^0^*, *C_t_^0^* (Ct also referred to as spontaneous torsion). C) i-iv: Predicted architectures of M1 polymer shape and membrane protrusion depending on different combinations of mechanical properties listed below each scenario.

**Table 1:**

| *Variable* | *Parameter* | *Value* | *Source* |
| --- | --- | --- | --- |
| **M1** |  |  |  |
| a, w, h | M1 Monomer length, width, height | 2.8 nm, 5 nm, 3.6 nm | (19)/ This study |
| $\Gamma_u, \Gamma_v, \Gamma_t$ | M1 strand Rigidities | $14\text{-}57 \times 10^{-27} \text{ Nm}^2$ | $\Gamma_u = Y I$ with<br>$I = (h^3w + w^3h)/12$<br>This study (Methods) |
| $2\pi b_{n=3}$ | Helix pitch (for n=3) | 10.8 nm | (19)/ This study |
| n | Number of helix starts | -2 to 6, 3 on average | (19) |
| D | Diffusion coefficient of M1 monomers | $\sim 10^{-11} \text{ m}^2\text{s}^{-1}$ | (29, 30) |
| Y | Young Modulus | Typically $10^9 \text{ Pa}$ for proteins | (34) |
| $\rho$ | M1 density per virion unit length | $\sim 12.5 \text{ nm}^{-1}$ | This study |
| $j$ | Diffusive M1 flux | | |
| $C_u, C_v, C_t$ | M1 strand curvatures | | |
| L | M1 filament length |  |  |
| f | Polymerization force |  |  |
| $\theta$ | M1 polymerization angle | | |
| **Membrane** |  |  |  |
| $\sigma$ | Membrane tension | Typically $10^{-5}$ to $10^{-3} \text{ N/m}$ | (41, 42) |
| $K_m$ | Membrane bending modulus | $4 \cdot 10^{-20} \text{ J}$ (lipid bilayer) to $10^{-16} \text{ J}$ (fully coated membrane) | (22, 43) |
| $\delta s$ | Change in membrane surface area | | |
| $R^0$ | Spontaneous membrane curvature | | |
| **Virus** |  |  |  |
| R | Virus tube radius | 24 nm | (19) |
| $v$ | Virus growth speed | | |
| S | Virus tube surface area |  |  |
| $L_{virus}$ | Virus length | | |

The deformation energy per unit length *L* of a M1 strand is:

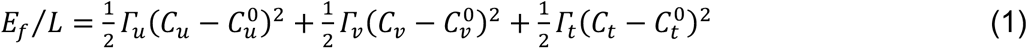

As M1 strands assemble on the membrane, several configurations are possible. M1 strands could form tubes of radius *R* underlined by either n juxtaposed helical strands (Fig. 2C case i) or straight strands running parallel to the filament axis (Fig. 2C case ii). Alternatively, strands could form a belt at the rim of the cell (Fig. 2C case iii). Case (ii) is the limit case of (i) for *θ → π/2*, in which *θ* is the angle of M1 strands relative to a plane orthogonal to the virus filament (table 1). Case (iii) is the limit of case (i) when *R* corresponds to the cell radius, and *θ → 0*. Therefore cases (ii) and (iii) are limit cases of case (i).

Finally, M1 strands could form a supercoiled helix (Fig. 2C case iv) in which C_u_ is non-zero. This would imply the formation of a helically bent virus filaments which is not observed experimentally, and this scenario can thus be excluded.

Hence, we will focus on the general case (i), for which we can write the deformation energy of both the coated membrane, and the helical M1 strands. The deformation energy per unit length of a coated membrane tube is:

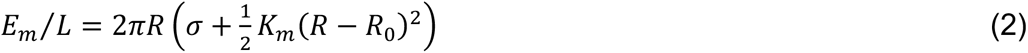

We further assume that each M1 strand takes the shape of a helix of constant pitch since a non-constant pitch would be energetically unfavorable. We define the pitch *b* such that 2π*b* is the periodicity of the M1 strand along the tube direction. Thus, with a tube radius R, we can write the deformation energy of helically-arranged M1 as:

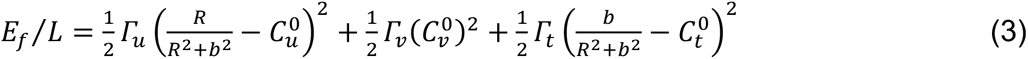

Since the second term is constant for a helix of constant pitch, it can be omitted when calculating the equilibrium shape by minimizing the elastic energy due to deformation. The other terms depend on the pitch and radius and their balance predicts how the shape is determined by the properties of the protein coated membrane (tension, rigidity, spontaneous curvature, equation 2) and on the properties of the M1 strands (stiffnesses and spontaneous curvatures, equation 3).

### Thermodynamic arguments predict packed helices of M1

From the established physical model, we can now predict the thermodynamically preferred organization of M1. High torsional rigidity *Γ_t_* of M1, associated to a spontaneous torsion *C_t_*>*0*, would favor unpacked helices, while membrane tension favors densely packed helices. We can compare the deformation energy for a virus-sized membrane tube (equation 2) to the deformation energy of M1 strands (equation 3) and predict the transition between packed and unpacked helices (Fig. 3A). Assuming *θ → 0,* we expect packed helices when the tension exceeds a critical value *σ*\*=Γ_t_C_t_ /2πR², corresponding to the helix elastic extension.

**Figure 3.**
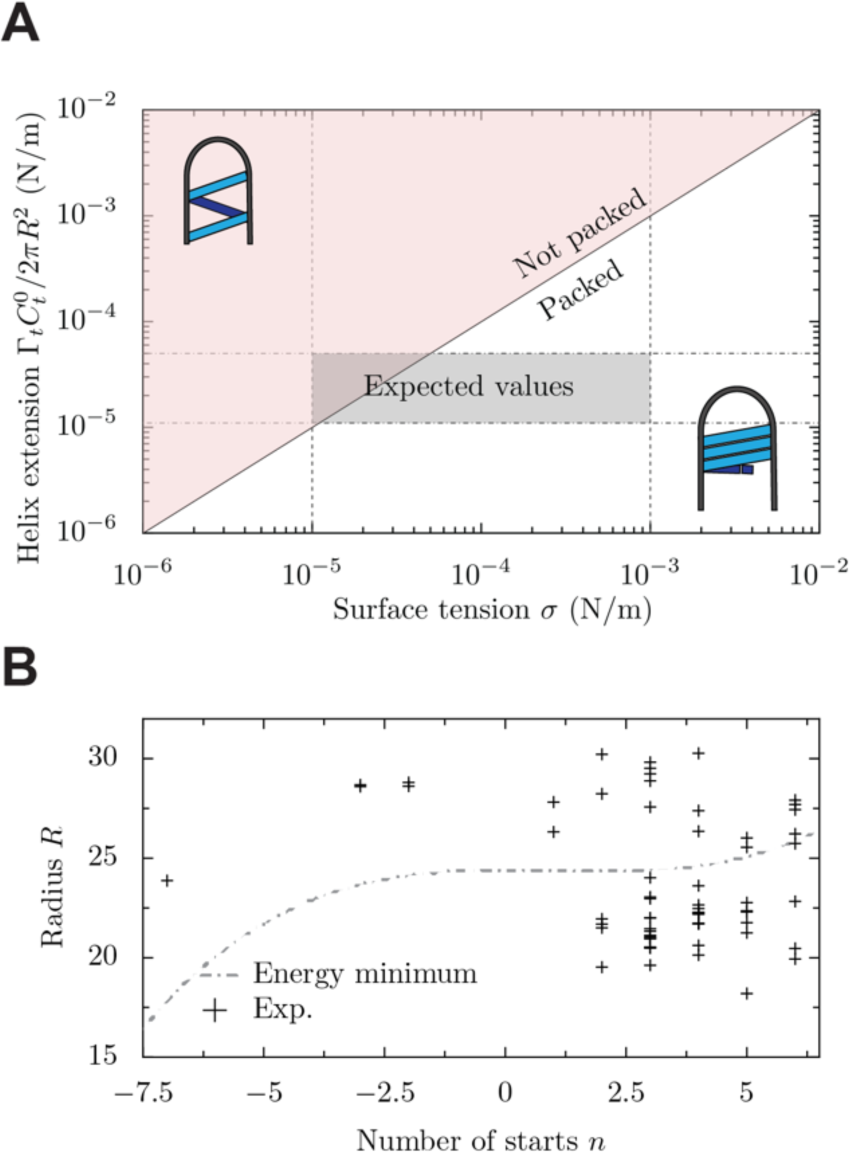
Thermodynamic predictions of M1 helical rise and radius. A) Theoretically predicted values of the M1 helical rise in dependence on membrane surface tension. Graphic insets illustrate the difference between scenarios of packed and unpacked helices. The expected physiological range of membrane tension and helical rise calculated based on documented values for sigma, R, C^0^ are marked. B) Experimental values for the number of helix starts and the tube radius, and the predicted relationship between the two variables by theoretical thermodynamic considerations. Experimental measurements do not suggest that helix start number is dependent on radius. The helix start number is likely to be kinetically controlled.

The typical elastic modulus of globular proteins (e.g. actin and tubulin (34), viral capsids (35), beta-barrel membrane proteins (36)) is in the order of the GPa (37). Assuming the elastic modulus of M1 to be in the same range, we can estimate the bending rigidities of an M1 strand *Γ_u_ , Γ_t_∼*14-57 10^⁻27^ *N m² (see methods).* For the plasma membrane, we expect a membrane tension σ between 10^-5^ and 10^-3^ *N/m (Table 1).* Our model suggests that, for this parameter range, helical M1 strands should be tightly packed (Fig. 3A), which is what we observed for M1 from experimental cryoET data (Fig. 1A).

### Virus filament radius largely depends on coated membrane properties, but not on membrane tension or helix start number

After having established what determines the helical parameters of M1, we sought to understand the mechanical determinants for the observed dimensions of the virus. We first considered whether the diameter of the virus filament should be dependent on membrane tension. Since the membrane tube is entirely covered by M1 strands, the increase of surface area due to a strand of length L is Δ*S= Lh* and thus independent of *R*. The energy cost of M1 strand extension due to membrane tension *σ* is Δ*E= σ* Δ*S* and therefore, changing the virus filament radius *R* does not alter the surface energy cost of the M1 strand, and as a result, virus filament radius is independent of *σ*.

We next considered whether the diameter of the virus filament should be related to the number of M1 strands. Because we predicted tight packing of M1 strands (Fig. 3A), we could further simplify our model by enforcing the resulting relationship between the number *n* of juxtaposed M1 strands (the number of helix starts), and the angle *θ* of the M1 strands, with *h* the height of a monomer:

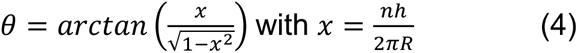

Thus, for a radius *R* and a strand number *n*, there is a single possible pitch 2π*b_n_:*

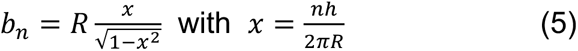

Thus, the helical pitch 2π *b_n_* depends on the radius R and the number of starts *n*. From equation 3, we know that the energy is a non-linear function of both *R* and *n,* and we expect a correlation between the radius and the number of starts. Since R>>b (as seen experimentally: b_max_=3.5 nm, R_min_=18.2 nm) the term depending strongly on *n* is the torsional term (dictated by *Γ_t_*). For a given radius, minimizing the energy yields an optimal number of starts:

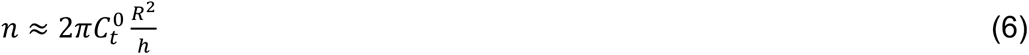

Alternatively, for a given number of starts, we can compute the optimal filament radius (see methods). Taking the limit of R being close to R_0_ *(i.e. R=R_0_(1+ε)* with *ε<<1)*, and assuming *Γ_t_*= *Γ_u_* for simplicity, we find to second order in *ε*:

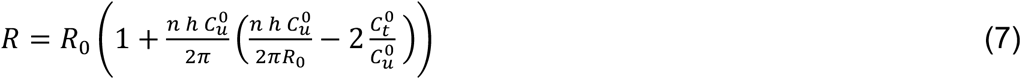

Thus, for a given *n*, *R* only marginally depends on *n, because R* is *C_u_^0^* plus a small correction of the order (*nhC_u_^0^/2π*)*^2^*. Consistent with this, experimentally, we find no correlation between R and *n* despite their variability (19), meaning we cannot thermodynamically explain the number of starts as a function of the radius, nor the radius as a function of the number of starts (Fig. 3B).

Recalling that the viral filament radius is independent of membrane tension, the radius should be set by the properties of the glycoprotein-coated membrane and/or by the M1 strand properties *Γ_u_* and *C_u_^0^*. We expect *Γ_u_*, *C_u_^0^* to be the same for all filaments – being internal properties of M1 proteins – while the elastic modulus of the membrane *K_m_*, and the spontaneous curvature of the membrane *R_0_* may both depend on the physiological state of the cell and might locally vary along the cell surface. The range of different radii that we observe thus suggests that physical properties of the coated membrane would differ between different cells, between different membrane regions in these cells, or at different stages of infection.

### A kinetic model predicts an optimal helix start number depending on membrane tension

Given that we find little difference in elastic energy between helices with variable number of starts, we next explored if the number of helix starts could be controlled by kinetics. This could be the case if as the tube begins to form, initially as a small membrane protrusion, the number of helix starts is determined by the number of M1 filaments nucleated. It is known that host proteins and viral RNP alter this nucleation process, but we could not find a difference in helix start numbers between virus and virus-like particles which do not carry a genome (19). Therefore, it seems unlikely that the viral RNPs would directly determine helix start number, although they could alter initiation kinetics in other ways.

Alternatively, the number of helix starts could influence the growth speed of the tube. Following our theoretical description of tube growth, we can estimate its growth speed as a function of the number of helix starts. Given a strand at an angle *θ*, adding a monomer increases the tube length by a value *a sin(θ)*, with *a* the length of a M1 monomer. While the viral tube has a radius R, the typical distance over which the membrane is deformed is rather *(K_m_/σ)^1/2^*; therefore, adding a monomer at the base of the tube should deform the membrane over an area δs ∼*a sin(θ) (K_m_/σ)^1/2^*. The energetic barrier to be overcome for polymerization is δe = *σ* δs, and the rate of monomer addition is thus *k_on_ = k_0_ exp(-δe /k_B_ T),* with *k_0_* the rate of monomer addition in the absence of membrane tension. Since *θ* is geometrically determined by the number *n* of starts, we can predict the growth speed *v_n_* of a tube:

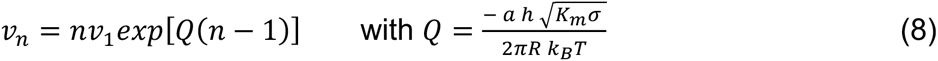

We find that the growth speed *v_n_* strongly depends on tension *σ* (Fig 4A) and that the maximum growth speed is attained for *n*=-1/Q* (Fig 4b). Plotting the predicted growth speed over the different number of helix starts for different tension values (Fig. 4A) reveals that for different membrane tension values, the curve peaks for different helix start numbers. In other words, depending on the membrane tension, a different helix start number yields the fastest growth. If we plot this optimal helix start number for a range of membrane tensions (Fig. 4B), we find that helix start numbers between 1 and 6 are most efficient for membrane tension values in the range 10^-4^ N/m to 3.8 10^-3^ N/m which lies within the physiological range of membrane tension values observed for mammalian cells (10^-5^ – 10^-3^ N/m ). We find that n=3, the helix start number which is most frequently observed, would lead to optimal growth speed for a tension of *σ* = 4.18 10^-4^ N/m *(table 1*) (Fig. 4).

**Figure 4.**
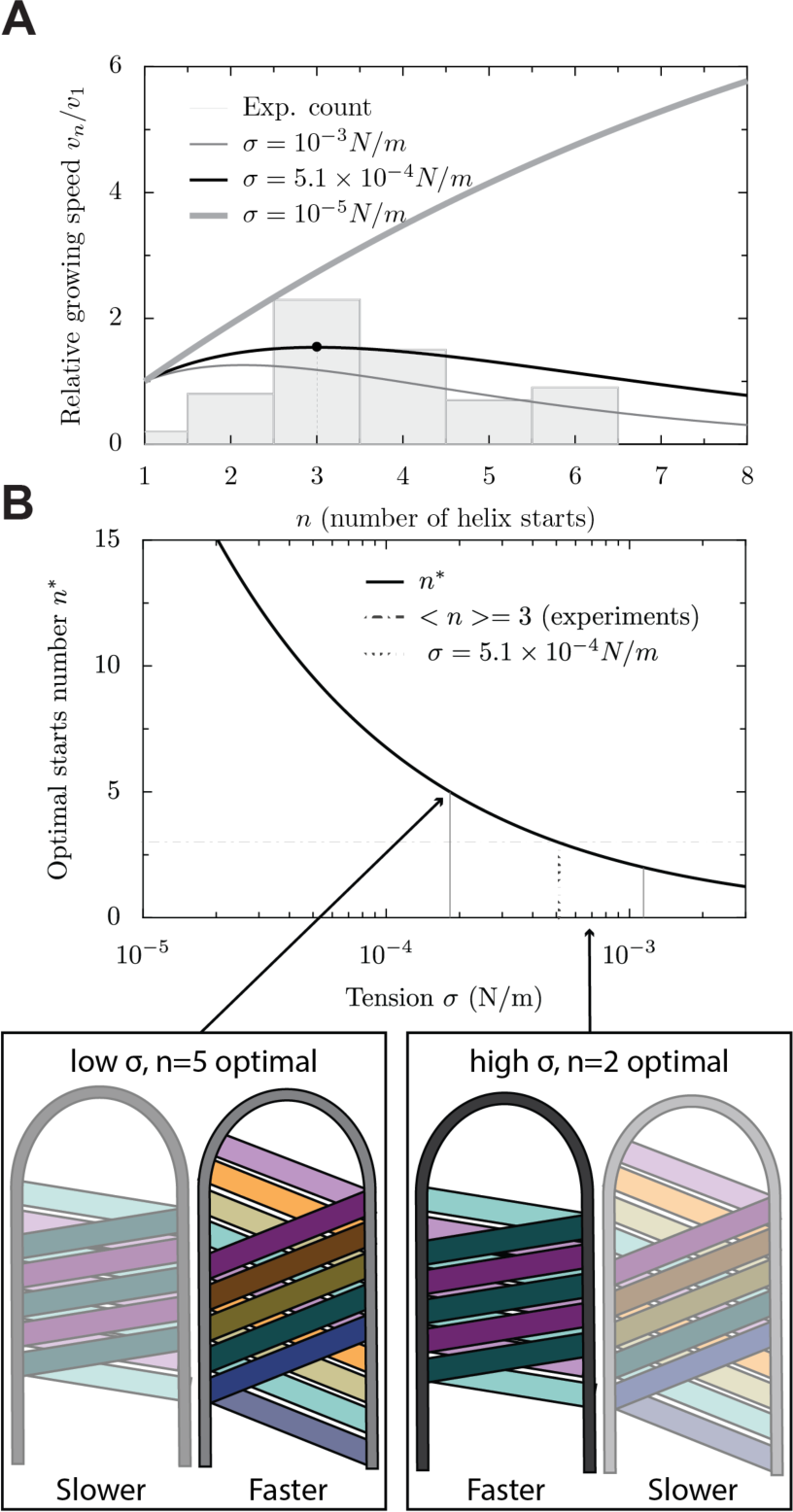
Predictions for kinetic control of M1 helix start number. A) Predicted filament growth speed plotted over the number helix starts. Curves are shown for three different values of membrane tension. The experimentally determined distribution of helix starts is plotted as a histogram. B) The optimal helix start number is plotted over membrane tension and ideal helix start numbers for different membrane tension regimes are marked. Schematics illustrate potential scenarios. Left: low membrane tension, more helix starts needed for optimal growth rate. Right: higher membrane tension, fewer helix starts needed for optimal growth rate.

### M1 polymerisation is sufficient to overcome membrane deformation resistance

Finally, we used our model in combination with experimentally determined values for virus and M1 dimensions to ask if M1 polymerization would generate enough energy to allow for virus tube formation. Structural and cryoET data allow us to estimate the dimensions of the virus as well as the numbers and dimensions of M1 strands, M1 monomers and the buried surface area between M1 monomers (24, 38) (Fig 2A, table 1). By feeding those values into our theoretical model, we can calculate the specific energy generated and required for M1 polymerization and virus tube formation respectively. From the previously determined structures of M1 we estimated the *δG* for the M1-M1 interface along the linear polymer to be 24 kcal/mol (19). Corresponding to ∼40k_B_T, this is an upper limit for the polymerization free energy of M1 strands. As a comparison the typical polymerization energy of actin is 1—10 k_B_T. The energy cost of tube formation due to membrane tension is Δ*E_1_ = σ* Δ*S_1_* per monomer, where Δ*S*_1_=10.1 nm^2^ is the surface area of one monomer on the membrane (M1 height h=3.6 nm and length a=2.8 nm (Fig. 2A, table 1)) and *σ* is the membrane tension. This is at most 2.5 k_B_T, and thus we estimate that M1-M1 association energy is indeed able to overcome the cost of surface tension in membrane tube extrusion.

## Discussion

M1, the most abundant protein in the virion, has been previously identified to play a major role in influenza virus assembly. M1 forms tightly packed linear strands arranging into helical arrays. Here we have used experimental data on M1 structure, M1 arrangement and virion size and morphology from cryoET and cryoEM, within a physical model to extract further insights into influenza virus assembly.

Virions can contain a variable number of interleaved M1 strands, and these strands form predominantly right-handed, but in some cases left-handed helices. Except for a minor subpopulation of virions which contain an additional protein layer, our analysis of the polarity of the M1 strands and the handedness of the helices reveals that they are oriented such that a free M1 N-terminal domain faces the cytosol. Assembly of M1 therefore has a defined directionality. We estimated the rate of M1 diffusion through the growing filament, and, with a concentration of M1 above 1 µM, this speed is compatible with the reported M1 polymerization rate. However, because one might expect that the speed of virion extrusion would provide a selection advantage, we favor a model where M1 assembles at the virion base, as this avoids the need of M1 diffusing through the tube, where diffusion might be hindered, and similarly avoids hindrance by the RNPs at the virion tip.

For assembly to proceed at the base of the virion, M1 monomers would be added to the growing polymer such that their CTD, which is unfolded in solution, binds the last unbound NTD, located at the rear end of the virus, and folds. It is unknown whether folding of the CTD is induced prior to interaction with the NTD, for example through allosteric effects of newly formed NTD-NTD interactions or NTD-membrane interactions, or whether the first interaction between the solution monomer and the growing strand is induced via folding of the CTD. Folding of the CTD carries an entropic cost, while formation of the large hydrophobic interface with the NTD releases energy, up to 40 k_B_T. While only a fraction of this energy may be released after folding of the CTD, a mere 2.5 k_B_T of polymerization free energy would be sufficient to overcome membrane tension and extend a filament, mostly because of the shallow angle of M1 strands with respect to the filament. We can therefore conclude that M1 strand polymerization provides sufficient energy to extend and protrude the growing virion.

Typically, in assembled influenza virions, multiple strands of M1 are arranged as a densely packed helix. This dense packing is predicted by our model, from the interplay between membrane tension and the deformability of polymerizing strands – it does not require any specific protein-protein interactions between M1 strands to form. This observation is consistent with the “slippery” nature of the inter-strand interactions that we have previously observed in the *in situ* M1 structure (19, 38).

Virion radius is likewise predicted by our model to be controlled thermodynamically based on a competition between the properties of M1 and the glycoprotein-coated viral membrane. Accordingly, the observed variability of radius in virus particles is expected to be due to variability in the coated membrane protein and lipid composition and properties at different places and stages of assembly. This may also include variable HA/NA protein rations on the virion surface (32).

Between one and seven parallel M1 strands are observed within virions. No strong dependence of the number of strands on virion radius was observed. This suggests a kinetic control of the number of strands likely based upon the number of nucleation events. Contribution and control of nucleation could come from additional viral or cellular components such as the vRNPs. We note however, that while vRNPs are present in the majority of viral particles they are absent in VLPs – while the number of M1 strands is comparable between VLPs and virions (19). This suggests that vRNPs are unlikely to regulate M1 strands number either because they do not contribute to the nucleation process or more likely because their contribution to nucleation leaves M1 polymerization kinetics unaffected.

The most common number of parallel strands in virions is 3 while 1–7 strands have been observed. Using a kinetic model of filament growth and reasonable estimates of the membrane tension, we calculated that this number of strands would yield the fastest filament growth for membrane tension values within the range expected for cellular plasma membranes. Hence, the frequency of M1 nucleation may have evolved to maximize virion growth speed.

Combining experimental data with a theoretical model has allowed us to consider the mechanics of influenza virus assembly. Our study supports a model where linear, polarised polymerization of M1 strands is the driving force for extension of filamentous influenza viruses. It suggests that nucleation kinetics have evolved to optimize the speed of filament growth for efficient membrane deformation under variable local conditions, thus increasing the robustness of virus assembly.

## Acknowledgements

We thank Hui Guo for helpful discussions. Funding was provided to J.A.G.B. by the Medical Research Council (MC_UP_1201/16) and the Max Planck Society and to S.D. by the CNRS “Emergence” program. The authors declare no competing interests.

---

**Supplementary Figure 1.**
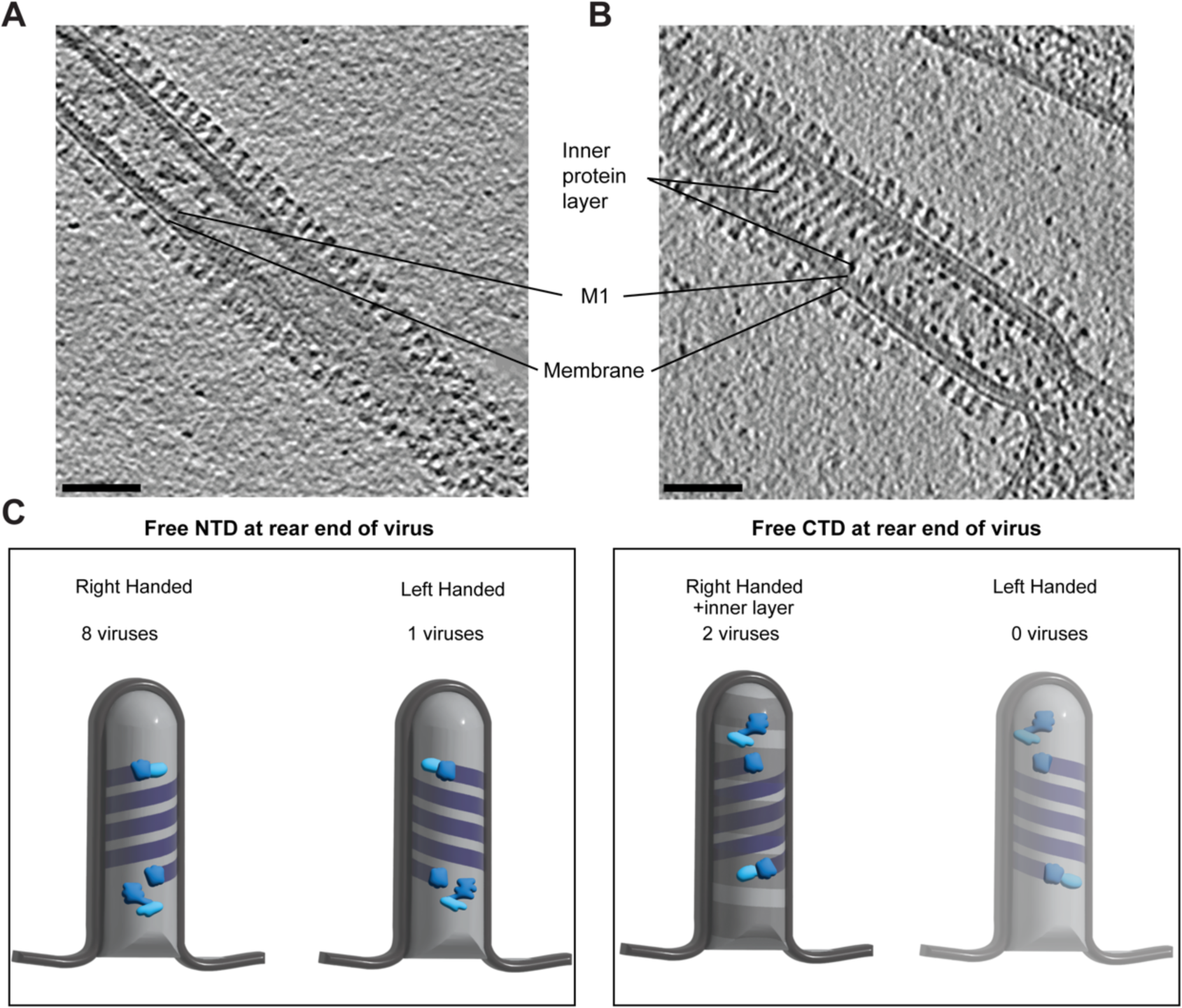
M1 orientation for M1 strands with different handedness and in the presence of an additional inner protein layer. A) Slice through a representative tomogram of an influenza A virus filament without an additional inner protein layer. B) Slice though a tomogram of an influenza A virus containing an additional protein layer at the inside of M1, which was present in 20 % of viruses. C) Illustrations of M1 orientation relative to M1 handedness and the presence or absence of an additional inner protein layer. Scale bars: 50 nm.

## Notes

### Competing Interest Statement

The authors have declared no competing interest.

